# Induction of mitophagy reverts age-associated decline of the hematopoietic and immune systems

**DOI:** 10.1101/2023.03.30.534925

**Authors:** Mukul Girotra, Yi-Hsuan Chiang, Melanie Charmoy, Pierpaolo Ginefra, Frederica Schyrr, Fabien Franco, Ping-Chih Ho, Olaia Naveiras, Johan Auwerx, Werner Held, Nicola Vannini

**Affiliations:** Department of Oncology, Ludwig Institute for Cancer Research, University of Lausanne; 1066 Epalinges, Switzerland; Department of Oncology, University of Lausanne; 1066 Epalinges, Switzerland; Laboratory of Regenerative Hematopoiesis, Department of Biomedical Sciences, University of Lausanne; and ISREC, School of Life Sciences, Ecole Polytechnique Fédérale de Lausanne (EPFL), Lausanne, Switzerland; Hematology Service, Department of Oncology, Centre Hospitalier Universitaire Vaudois (CHUV); Lausanne, Switzerland; Laboratory of Integrative and Systems Physiology, Institute of Bioengineering, Ecole Polytechnique Fédérale de Lausanne (EPFL); Lausanne, Switzerland

## Abstract

Aging compromises hematopoietic and immune system functions, making elderly individuals especially susceptible to hematopoietic failure, infections and tumor development and thus representing an important medical target for a broad range of diseases. During aging, hematopoietic stem cells (HSCs) lose their blood reconstitution capability and commit preferentially toward myeloid lineage (myeloid-bias). These processes are accompanied by an aberrant accumulation of mitochondria in HSCs. The administration of the mitophagy-inducer Urolithin-A re-establishes the correct mitochondrial homeostasis in HSCs and completely restores the blood reconstitution capability of “old” HSCs. Moreover, Urolithin-A supplemented food restores lymphoid compartments, boosts HSCs function and improves the immune response to viral infection in old mice. Altogether our results demonstrate that targeting mitophagy reverts aging phenotype in the hematopoietic and immune system.

## Main Text

Hematopoietic stem cells (HSCs) generate all the blood and immune cells throughout the entire life of an organism and they ensure the correct homeostasis between lymphoid and myeloid lineages. However, aging dramatically reduces HSC blood reconstitution capability and skews their fate toward myeloid lineages to the detriment of lymphoid cell production (*1, 2*). In fact, specific age-related dysfunction of the immune systems stems from the deteriorated HSC pool (*3, 4*). Thus it has been proposed that reverting HSC aging would improve both hematopoietic and immune functions (*5*). HSC aging is coupled with increased DNA damage due to an impairment of DNA-repair enzymes (*6, 7*) and epigenetic changes (*8, 9*). Interestingly, aging has also been directly linked to metabolic alterations in HSCs (*10*). These studies have led to the emerging concept that cellular metabolism operates as a potent regulator of the HSC pool (*11-13*), and more broadly, of stem cell function and fate (*10, 14, 15*). Defective autophagy and consequent accumulation of mitochondria in HSCs has been associated with aging processes (*16*), providing evidence that cellular quality control processes are crucial for the maintenance of a functional HSC pool both in homeostasis and in stress conditions (*17-19*). Even though HSCs are refractory to systemic rejuvenating approaches (*20*), nutritional interventions such as prolonged fasting and NAD boosting strategies have been demonstrated to partially restore HSC function in old mice (*21, 22*). In particular, NAD precursors have been shown to exert their effect by directly modifying HSC mitochondrial function (*18, 22*).

In this context, Urolithin-A (UroA), a natural compound belonging to the ellagitannin family (*23*), has been shown to improve metabolic fitness in human and rodents, and prolong life span in worms via the induction of mitophagy (*24-27*). Because HSCs have defective autophagy and accumulate mitochondria during aging (*16, 28*) (fig. S1), we tested the capacity of UroA to recover the functions of HSCs purified from >18 months old mice (“old HSCs”). Purified old HSCs (lin^-^ Sca1^+^ cKit^+^ CD48^-^ CD150^+^) cultured for three days in the presence of 20μM Uro-A were transplanted into lethally irradiated recipient mice and blood chimerism was assessed over a period of 24 weeks by exploiting the CD45.1/2 double congenic allelic system (fig. 1A). The comparison of the blood reconstitution capabilities revealed that exposure to UroA restored the reconstitution function of old HSCs to levels seen with HSCs purified from young mice (8-12 weeks; young HSCs) (fig. 1B). The functional restoration of old HSCs was further confirmed by an increased donor chimerism in primary and secondary lymphoid tissues, bone marrow and spleen respectively (fig. 1C). To ascertain the long-term HSC gain of function, bone marrow from primary recipient mice was transplanted into lethally irradiated secondary recipient mice and donor chimerism analysis was followed over a period of 20 weeks (fig. 1A). The UroA effect was maintained during secondary transplantation, where the UroA treated old HSCs sustained an identical blood reconstitution capability of young HSCs (fig. 1D), produced a comparable donor-derived chimerism in primary and secondary lymphoid tissues (fig. 1E, S2) and restored multi-potent stem and progenitor population (LKS: lin^-^ cKit^+^ Sca1^+^) in the bone marrow (fig. 1E).

**Fig. 1.**
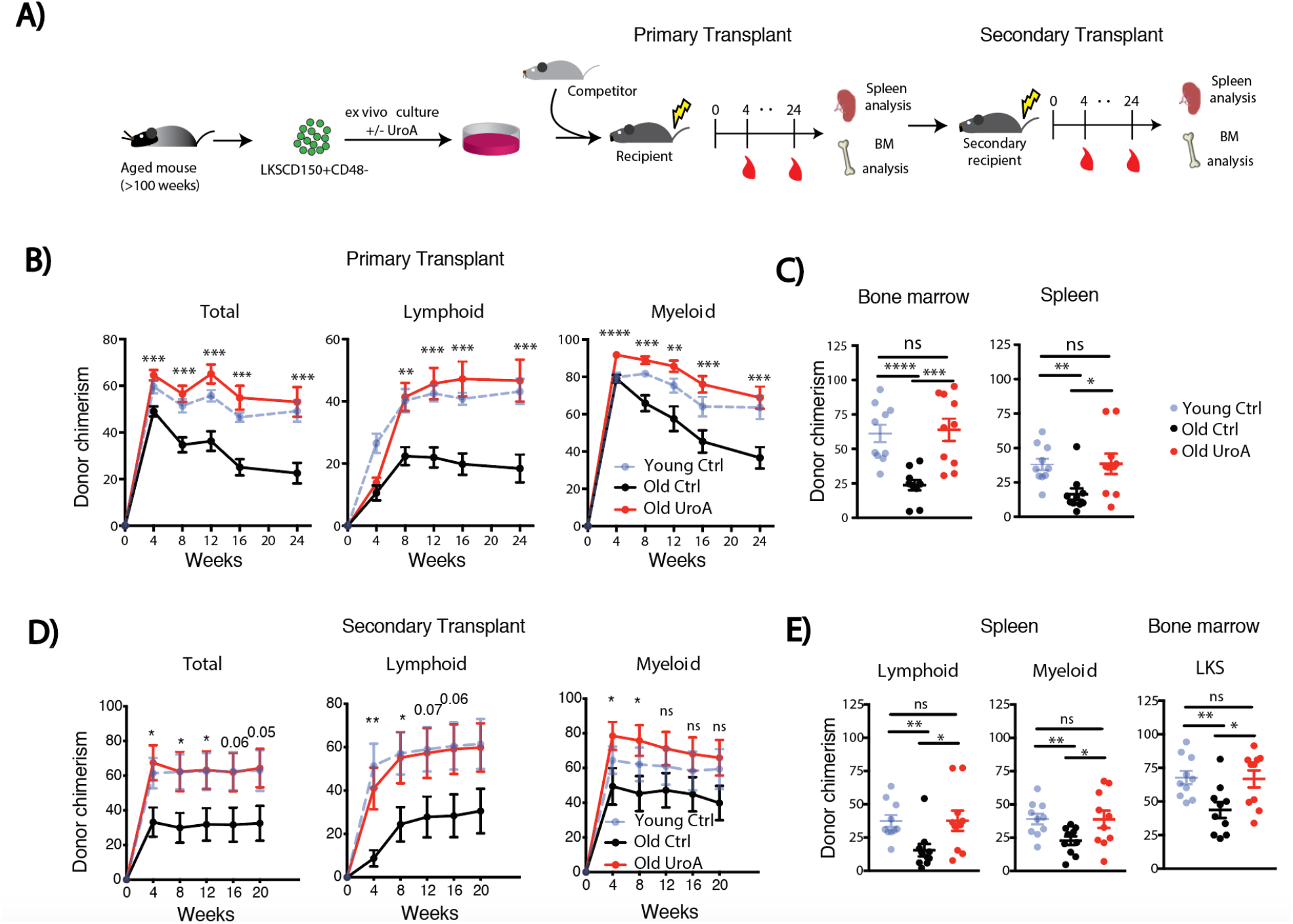
UroA *in vitro* treatment reverts age-associated defects in HSCs. (**A**) HSCs isolated from mouse bone marrow were cultured for three days in the presence of UroA and transplanted in lethally irradiated primary recipient together with bone marrow derived from competitor mice. At the end point the bone marrow of primary recipient was transplanted in lethally irradiated secondary recipient mice. (**B**) Blood donor chimerism analyses of primary transplant at indicated timepoints. (**C**) Spleen and Bone marrow analyses of donor chimerism at endpoint. (**D**) Blood donor chimerism analyses of secondary transplant at indicated timepoints. (**E**) Spleen analyses of donor chimerism at endpoint and analysis of donor-derived stem/multi-potent progenitor population LKS (Lin-cKit+Sca1+). (*n*=10; Student’s *t* test; * P ≤0.05, ** P ≤ 0.001, *** P≤0.001, **** P≤0.0001).

Similarly, UroA treatment improved hematopoietic stem and progenitor activity of human CD34^+^ cells purified from elderly patients, as evidenced by improved colony forming capacity (fig. S3). Taken together these data highlight that a short *in vitro* exposure (three days) to UroA restored the long-term blood reconstitution capability of old HSCs.

To study whether UroA recovered hematopoietic and immune functions when delivered systemically *in vivo*, we fed mice with an UroA enriched diet for a period of 4 months and characterized hematopoietic stem and progenitor pools and analyzed immune compartments (fig. 2A). The cellular composition of the blood, which was monitored monthly, was not different between UroA treated and untreated groups (fig. S4). However, analyses of hematopoietic stem and progenitor populations of the bone marrow (fig. S5A) revealed an expansion of hematopoietic stem cells and common lymphoid progenitor cells (CLPs: lin^-^ CD127^+^ Sca1^low^cKit^low^) and a reduction of megakaryocyte erythroid progenitor cells (MEPs: lin^-^ cKit^+^ Sca1^-^ CD16/32^-^ CD34^-^) (fig. 2B), while no changes were observed in common myeloid progenitor cells (CMPs: lin^-^ cKit^+^ Sca1^-^ CD16/32^-^ CD34^+^), granulocyte monocyte progenitor cells (GMPs: lin^-^ cKit^+^ Sca1^-^ CD16/32^+^ CD34^-^) and multipotent progenitor cells (MPPs: lin^-^ Sca1^+^ cKit^+^ CD48^-^ CD150^-^) (fig. S5B). Indeed, during aging CLPs are reduced while megakaryocyte progenitors expand (*29*), thus UroA supplementation reverted aging associated features of the hematopoietic system. Interestingly, CLPs expansion correlated with a partial expansion of lymphocytes in the spleen (fig. 2C). To test bone marrow functionality of old mice supplemented with UroA, we transplanted whole bone marrow into lethally irradiated mice (fig. 2A). Donor chimerism analyses revealed that bone marrow derived from UroA treated old mice have higher reconstitution capacity compared to the untreated group (fig. 2D), which was maintained in secondary transplantation (fig. S5C).

**Fig. 2.**
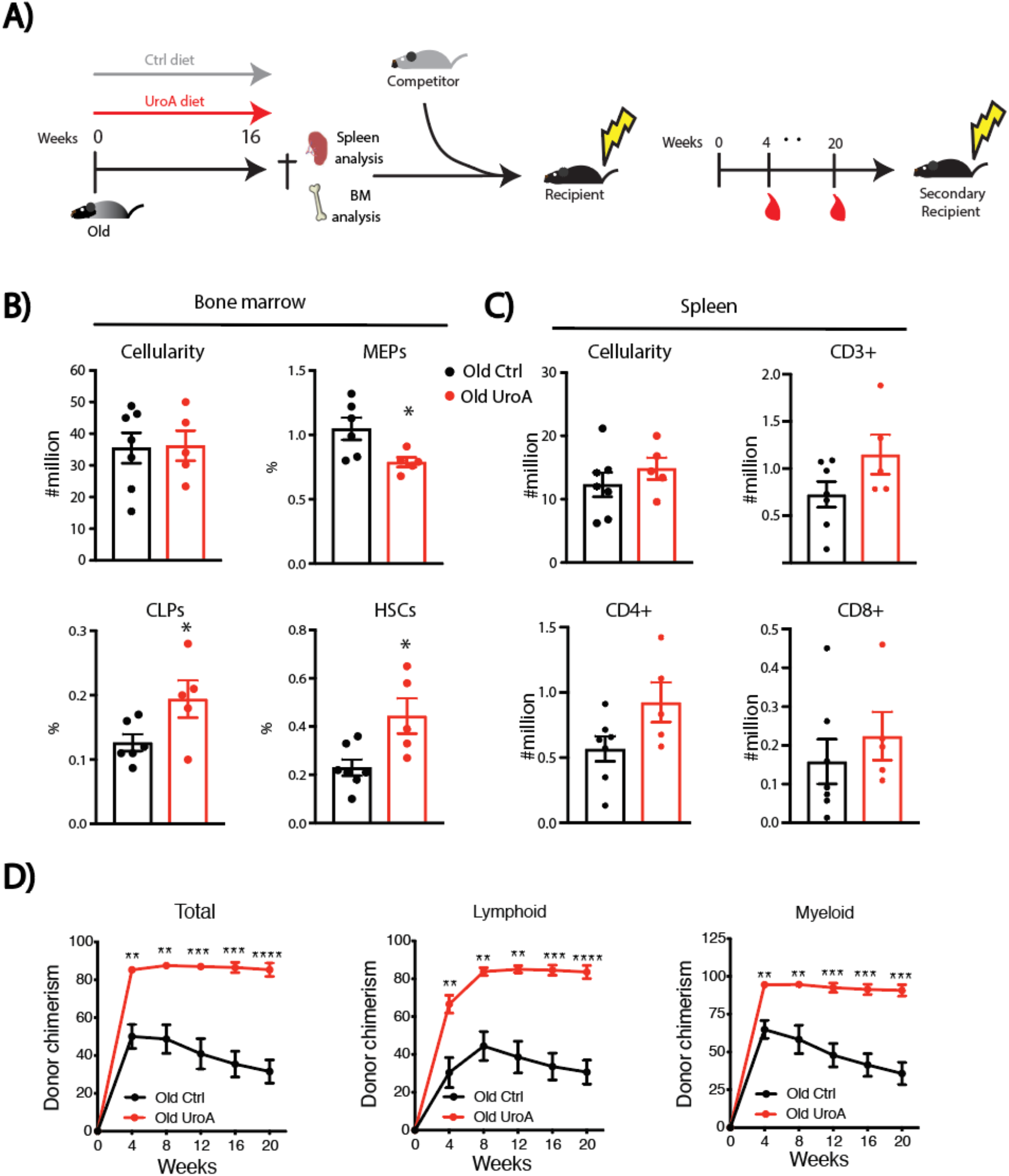
UroA oral supplementation rejuvenates the hematopoietic system of old mice. (**A**) old mice were fed with control or UroA supplemented diet (50mg/kg/day) for 16 weeks. Mice were sacrificed to analyse the spleen and BM. A part of the BM was transplanted in lethally irradiated primary and secondary recipients. (**B**) Analysis of different stem and progenitor populations of the BM by flow cytometry. (**C**) Analysis of different T cell compartments in the spleen by flow cytometry (*n*=5; Student’s *t* test; * P≤0.05, ** P≤0.001, *** P≤0.001). (**D**) Blood donor chimerism analyses of primary transplant at indicated timepoints (*n*=5; Student’s *t* test; * P≤0.05, ** P≤0.001, *** P≤0.001, **** P≤0.0001).

These data indicate that UroA dietary supplementation boosts hematopoietic stem and progenitor functions and restore the CLP population in old mice.

To evaluate whether the boost of hematopoietic capacity would also reflect an improvement in immune functions, old mice supplemented for 8 weeks with an UroA enriched diet were infected with lymphocytic choriomeningitis virus (LCMV) Armstrong strain, which results in an acute resolved infection depending on a CD8^+^ T cell response. The specific immune response against the virus was assessed at day 8 post-infection (fig. 3A, B). Aging strongly reduces the abundance of virus-specific CD8^+^ T cells and the signaling via IL-7 pathway, increases the expression of co-inhibitory receptors such as PD1 and LAG3 and impairs T cell functionality (*30, 31*). UroA supplementation did lead to the expansion of virus specific CD8^+^ T cells recognizing the virus specific antigens np396 (fig. 3C) but not to an improved production of the inflammatory cytokines INFγ, TNFα and IL-2 (fig. S6A). However, for both antigens UroA supplementation restored the expression of IL7 receptor (CD127) on memory precursor cells(fig. 3D) and improved terminal differentiation of old CD8^+^ T cells as indicated by increased KLRG1 (fig. 3E) and Granzyme B expression by virus specific CD8^+^ T cells (fig. 3F), indicating improved cytolytic effector differentiation. Moreover, virus-specific CD8+ T cells expressed reduced levels of the inhibitory PD-1 and LAG3 receptors (fig.S6B). Consistent with increased effector differentiation and function, UroA supplemented old mice showed improved virus control (fig. 3G).

**Fig. 3.**
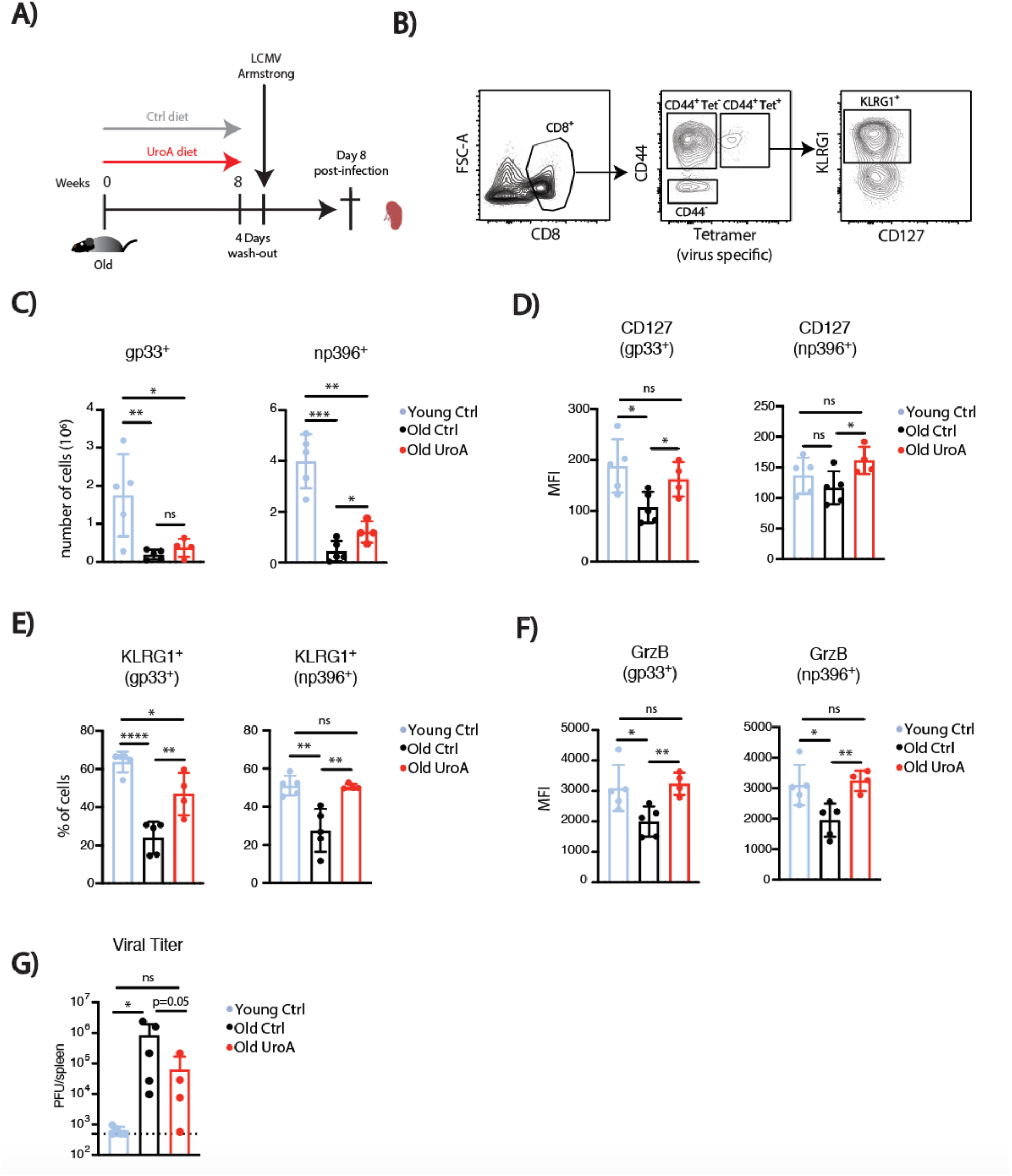
UroA supplementation improves the response to viral infection of old mice. (**A**) old mice were fed with control or UroA supplemented diet (50mg/kg/day) for 8 weeks. Mice were sacrificed to analyse the spleen (**B**) Gating strategy for virus-specific CD8 T cells and their immune profile. (**C**) Quantification of activated (CD44^+^) virus-specific CD8 T cells (Tet^+^: gp33^+^ or np396^+^). (**D**) Expression analysis of IL-7 receptor (CD127) in CD44^+^gp33^+^ and CD44^+^ np396. (**E**) Analysis of the differentiation marker KLRG1 in CD44^+^gp33^+^ and CD44^+^ np396^++^. (**F**) Analysis of granzyme B expression in stimulated CD8+ T cells. (**G**) Quantification of viral titer in the spleen (*n*=5; Student’s *t* test; * P ≤ 0.05, ** P ≤ 0.001, *** P ≤ 0.001).

To assess whether the UroA effect was exerted through mitophagy induction, we tested the capacity of UroA to induce mitophagy in HSCs. HSCs purified from mitophagy reporter mice mito-QC were cultured for 3 days in the presence of 20μM UroA and mitophagy induction was measured by flow cytometry. Mito-QC mice express a mCherry-GFP tandem protein targeted to the mitochondrial outer membrane. When mitochondria undergo mitophagy, the acidic environment of lysosomes quenches the GFP but not the mCherry fluorescence and the induction of mitophagy can be measured based on the shift on the GFP signal (*18, 32, 33*).

Indeed, UroA treatment induced mitophagy in HSCs (fig. 4A). Importantly induction of mitophagy was accompanied by an improved spare respiratory capacity (%SRC) (fig. 4B), indicating improved mitochondrial performance. Since HSCs accumulate mitochondria due to autophagy defects (fig. S1), and have increased expression of CD150 during aging (fig. S7) (*16, 28*), we investigated the capacity of UroA to revert these aging phenotypes in old HSCs. UroA reduced CD150 expression and mitochondrial mass in purified old HSCs (fig. 4 C, D). Interestingly, old CD150^high^ HSCs, which have been shown to have a marked myeloid-bias and aging phenotype (*28*), have higher mitochondrial content as compared to CD150^low^ HSCs. Treatment with UroA comparably decreased the mitochondrial content in both CD150^high^ and CD150^low^ HSCs (fig. S1B). These data indicate that UroA treatment re-establishes mitochondrial homeostasis and fitness *via* mitophagy induction.

**Fig. 4.**
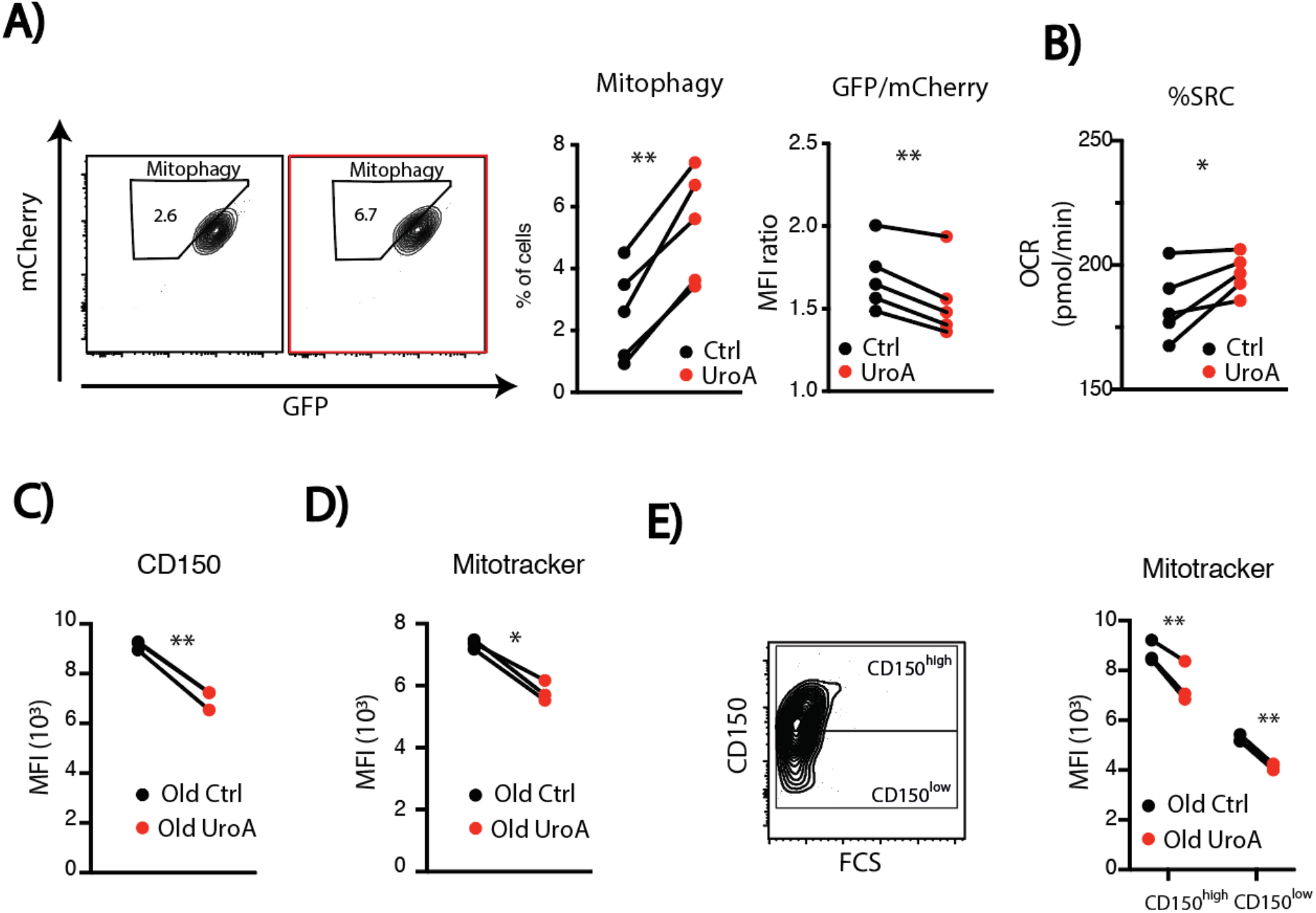
UroA induces mitophagy and restores mitochondrial fitness. (**A**) Analysis of mitophagy induction measured as percentage of cells in the mitophagy gate (left) and mean fluorescence intensity (MFI) ratio between GFP and mCherry signals. (**B**) Analysis of spare respiratory capacity (SRC) measured *via* oxygen consumption rate (OCR) in Lin-cKit+ hematopoietic stem/progenitor population. (*n*=5; Student’s *t* test; * P≤0.05, ** P ≤0.001, *** P ≤ 0.001) (**C**) CD150 expression and (**D**) mitochondrial mass (mitotracker) in old HSCs treated (Old UroA) or not treated with UroA (Old Ctrl), values are expressed as MFI. (**E**) Measurement of mitochondrial mass by mitotracker green staining in CD150^high^ and CD150^low^ HSCs expressed as MFI and evaluation of UroA treatment effect. (*n*=3; Student’s *t* test; * P ≤ 0.05, ** P ≤ 0.001, *** P ≤ 0.001).

The loss of cellular quality control processes during aging leads to the accumulation of defective cellular machineries and to a rapid deterioration of cellular function (*34*). Our results demonstrate that treatment with the mitophagy inducer, UroA, is capable to revert the metabolic defects in old HSCs and rejuvenate hematopoietic and immune system functions providing further evidence on the important interplay between cellular metabolism and aging-related cellular dysfunction.

## Acknowledgments

We thanks prof. George Coukos (LICR, UNIL, CHUV) for critical feedback on the study. We thank Francis Derouet (UNIL) for animal care. Flow cytometry analysis and sorting was performed at Flow Cytometry Facility at UNIL with the help of Romain Bedel, Francisco Sala de Oyanguren, Kevin Blackney and Anne Wilson. We thank Nestlé Health Science for their suggestions on the study.

## Author contributions

MG, NV conceived the ideas, designed the experiments and analyzed the results. NV wrote the manuscript. MG, YHC, NV and FS performed hematopoietic stem cell experiments. WH and CH performed and interpreted the infection model experiment. PG evaluated the impact of UroA dietary supplementation on secondary lymphoid tissues, performed and interpreted cell respiration analyses. JA and ON conceived ideas and provided key reagents and samples. FF and PCH provided animal model and helped on data interpretation. All authors edited and reviewed the final manuscript.

## Data and materials availability

All data and materials used in the analysis are available to any researcher for purposes of reproducing or extending the analysis.

## Supplementary Materials

Materials and Methods

Figs. S1 to S7

References 31-34

