## supplementary data and material for "Induction of mitophagy reverts age-associated decline of the hematopoietic and immune systems"

**This PDF file includes:**

Materials and Methods
Figs. S1 to S7
References 31-34

### Materials and Methods

#### Bone marrow extraction and cell sorting

Flow cytometry analysis was performed on freshly isolated bone marrow (BM) from C57Bl6 mice (young: 8 weeks old; old > 80 weeks). BM was extracted from crushed femur, tibia and pelvis. Cells suspension was filtered through 70µm cell strainer and erythroid cells were eliminated by incubation with red blood cells lysis buffer (eBioscences). Isolation and stains were performed in ice-cold PBS 1mM EDTA. Lineage positive cells were then removed with a magnetic lineage depletion kit (BD biosciences). Cells suspension were then stained with specific antibodies for progenitor and stem cell compartments and sorted on BD FACS Aria III into 1.5ml Eppendorf tubes. The hematopoietic stem cell (HSC) compartment was identified and sorted with the following cell surface phenotype $Lin^-$ $Ckit^+$ $Sca1^+$ (LKS) $CD150^+$ $CD48^-$.

#### Antibodies

The following antibodies were used in this study: rat mAbs against Stretavidin- Texas red, cKit-PeCy7 (2B8), Sca1-APC (D7), CD150- PE (TC-15-12F12.2), CD48-PB (HM48-1), CD45.2-PB (104), CD45.1-FITC (A20), Gr1-APC (RB6-8C5), F4/80-APC (BM8), CD19-PE (6D5), CD3-PE (17A2), CD16/CD32 (2.4G2). The antibodies were purchased from Biolegend, eBiosciences and BD. A mixture of biotinylated mAbs against CD3, CD11b, CD45R/B220, Ly-6G, Ly-6C and TER-119 was used as lineage marker ("lineage cocktail") and was purchased from BD. Human specific antibodies were: hCD56 (NCAM16.2), hCD16 (3G8), hCD45 (HI30), hCD19 (HIB19), hCD4 (RPA-T4), hCD3 (SK7), hCD14 (M5E2), hCD8b (SIDI8BEE), hCD34 (8G12), hCD38 (HB-7) and were either from eBioscience or BD. DAPI staining was used for live/dead cell discrimination.
For viral infection analyses the following antiobodies were used: CD44-PB (IM781), KLRG1-APC (2F1), CD127-BV786 (A7R34), PD-1-PE-Cy7 (RMP1-30), LAG3-PercPCy5.5 (eBioC9B7W), GrzB-PB (GB11), IFNγ-PercPCy5.5 (XMG1.2), TNFα-PE-Cy7 (MP6-XT22), IL2-APC (JES6-5H4).

#### Mouse HSC/progenitor and human CD34+ cells culture

Murine HSCs were sorted in 1.5ml eppendorf tubes. Cells were seeded in 96-well round bottom plates and cultured in Stemline II (SIGMA) supplemented with 100ng/ml SCF (R&D) and 2ng/ml Flt3 (R&D). UroA (Sigma) was added at indicated concentrations (dissolved in DMSO), equal quantity of DMSO was added to the control wells. All cultures were maintained at 5% $CO_2$ at 37$^o$ C.
Cryopreserved CD34+ cells isolated from from hip replacement orthopedic patients were thawed and cultured in StemSpan (Stem cell tech) media supplemented with hSCF (100ng/ml), hFLT3L (100ng/ml), hTPO (50ng/ml), hLDLP (10ug/ml) and different concentrations of UroA (dissolved in DMSO) were added, equal amount of DMSO was added to the control wells. For longer culture periods half of the media was replenished every 2$^{nd}$ or 3$^{rd}$ day. All cultures were maintained at 5% $CO_2$ at 37$^o$ C.

#### Analysis of mitochondrial mass by flow cytometry

Post culture cells were incubated at 37°C for 45 minutes with 100nM Mitotracker green. Cells were then washed with FACS buffer and analyzed by flow cytometry on BD LSR II.

**DNA extraction and mtDNA/ nuDNA estimation**

Progeny of 1000 cells were collected at the end of the culture period and DNA was isolated using DNeasy kit (Qiagen) according to the manufacturer's instruction. QPCR was carried out to estimate the relative values for mitochondrial (Cox 1) and nuclear (NDUFA2) gene to estimate mt/nu DNA ratio.

**Murine Bone marrow transplantations**

C57Bl/6 Ly5.2 adult mice (8-12-week-old) were lethally irradiated with a total 8.5Gy dose in a X-ray irradiator (RS-2000, RAD source) 24h before transplant. The dose was split in two doses of 4.25Gy separated by a 4-6 hour interval. Mice were injected with 2000 donor cells post culture derived from C57Bl/6 Ly5.1 mice (young ot old) and 150,000 competitor cells derived from C57Bl/6 Ly5.1/5.2 mice, via tail-vein injection. Peripheral blood was collected every 4 weeks to determine the percentage of chimerism by FACS analysis. Spleen and bone marrow were analysed at the endpoints. For secondary transplants, adult mice (8-12-week-old) were lethally irradiated with the same dose 24h before transplant and each mouse was injected with 3 million whole bone marrow cells from a donor mouse.

**CFU assay**

CFU assay was carried out using methocult H4434 (Stem cell tech) as per manufacturer instructions. 1000 cells from each well were plated in duplicates. Colonies were counted 15 days post plating using Stem Vision (Stem cell tech).

**Food preparation for in vivo feeding**

UroA diet was prepared by mixing custom synthesized (by Novalix) UroA dissolved in DMSO with 2916 powder diet (Charles River) and air-dried into pellets in sterile conditions. The UroA dosage in the mix was calculated by considering the average mouse food intake per day (5g food/ mouse/ day) for a calculated intake of 50mg UroA /mouse/day). Equivalent amount of DMSO was added in the control food. Weekly food consumption was monitored in all cages and no difference was found in the feeding behaviour of mice upon UroA supplementation in the diet.

**Viral infections and staining**

Mice were infected intraperitoneally (i.p.) with $2x10^5$ plaque forming units (pfu) of LCMV 53b Armstrong (Arm) strain. To determine viral titers, spleens from LCMV-infected mice were 'shock frozen'. Diluted spleen suspensions were then used to infect Vero cells, and viral titers were determined by an LCMV focus-forming assay, as described elsewhere (Battegay et al.,1991).

Cell suspensions from the spleen were obtained by mashing through a 40µM nylon cell strainer, followed by red blood cells lysis using ACK buffer. Surface staining was performed with mAbs for 20 min at 4°C in PBS supplemented with 2% FCS (FACS buffer).

For tetramer staining, cell suspensions were incubated with anti-CD16/32 (2.4G2) hybridoma supernatant before staining for 90min at 4°C with APC-conjugated MHC-I tetramers.
For intranuclear staining, cells were surface stained before fixation and permeabilization using the Foxp3 transcription factor staining kit (eBioscience: Cat. No. 00-5523) followed by intranuclear staining in Permeabilization buffer 1x (Perm buffer).
For the detection of cytokine production, splenocytes were re-stimulated in vitro with LCMV gp33- 41 (gp33) or np396-404 (1μM) peptide for 5h in the presence of Brefeldin A (5μg/ml) for the last 4.5h. Cells were then stained at the surface before fixation and permeabilization (using eBioscience kit: Cat. No. 88- 8824) followed by intracellular staining in 1x Perm buffer.

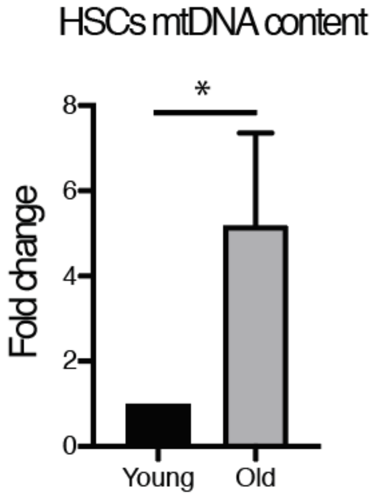


**Fig. S1. Old HSCs have higher mitochondria content.** Mitochondrial DNA (mtDNA) quantification by QPCR expressed as fold change compare to young HSCs (*n* = 4; Student's *t* test; * P ≤ 0.05, ** P ≤ 0.001, *** P ≤ 0.001).

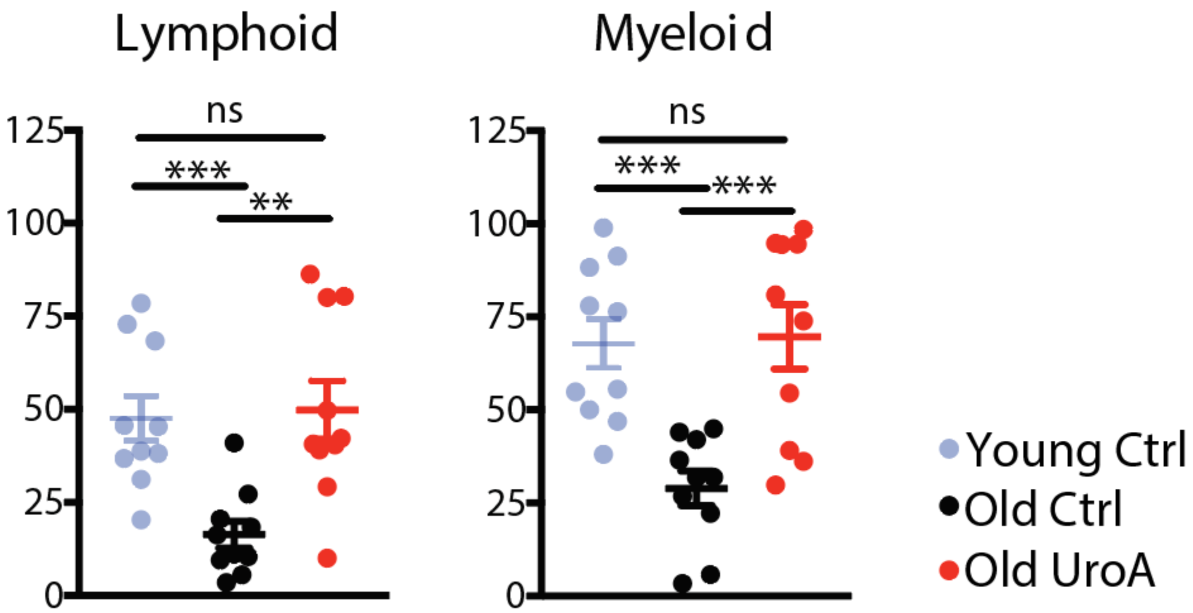


**Fig. S2. Donor chimerism analysis in the bone marrow.** Analysis of donor derived contribution for lymphoid and myeloid lineages in bone marrow. ( *n*=10; Student's *t* test; * P ≤ 0.05, ** P ≤ 0.001, *** P ≤ 0.001, **** P ≤ 0.0001).

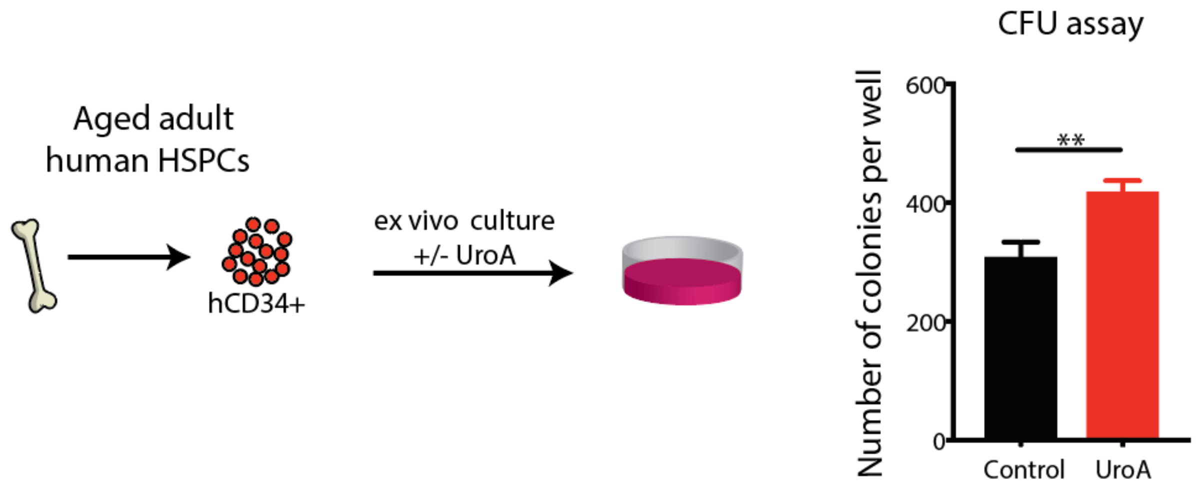


**Fig. S3. Colony forming capacity (CFU assay) of aged human CD34$^{+}$**. CD34$^{+}$ cells derived from elderly patients were treated for 3 days with UroA, and colony formations was measured at 15 days post seeding (*n*=3. Student's *t* test; * P ≤ 0.05, ** P ≤ 0.001, *** P ≤ 0.001).

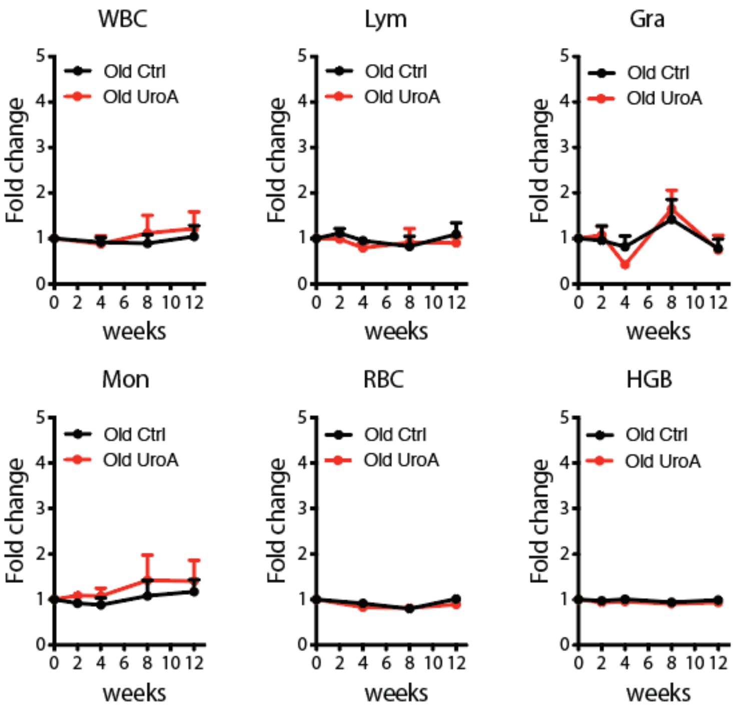


**Fig. S4. Peripheral blood analyses of UroA fed mice.** Peripheral blood cell analyses of old mice fed with ctrl (old ctrl) or UroA enriched diet (old UroA). Analyses were performed with blood cell counter and normalized to day 0 (WBC: white blood cell; Lym: Lymphocites; Gra: Granulocytes; Mon: Monicytes; RBC: red blood cells; HGB: hemoglobin) (*n*=8).

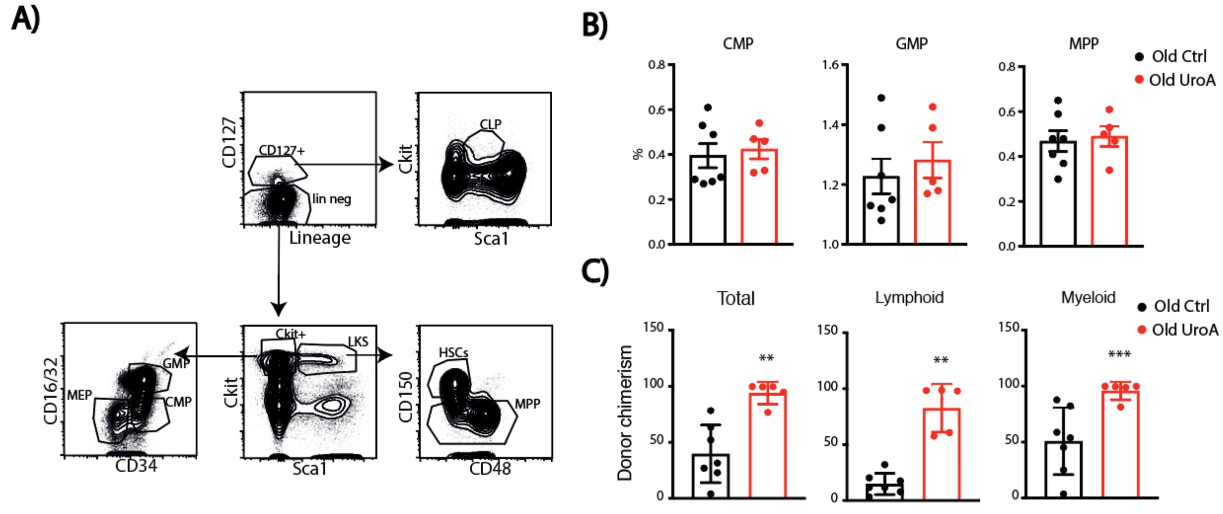


**Fig. S5. Hematopoietic stem and progenitor analyses of old mice supplemented with UroA.** (**A**) Gating strategy to identify various stem and progenitor populations from the BM for endpoint analysis. HSCs: Lineage$^{-}$cKit$^{+}$Sca1$^{+}$ (LKS) CD150$^{+}$CD48$^{-}$; Multipotent progenitors (MPPs): LKS CD150$^{-}$, Common lymphoid progenitor (CLP): CD127$^{+}$ cKit$^{low}$ Sca1$^{low}$; Committed progenitors (cKit$^{+}$) include common myeloid progenitors (CMPs: Lineage$^{-}$ cKit$^{+}$Sca1$^{-}$(KLS$^{-}$) FcRlowCD34$^{+}$), granulocyte–macrophage progenitors (GMPs: KLS-FcRlowCD34+), and megakaryocyte-erythroid progenitors (MEPs: KLS$^{-}$FcR$^{-}$CD34$^{-}$). (**B**) Analysis of different stem and progenitor populations of the BM in response to UroA feeding ($n$=5). (**C**) Donor chimerism analysis of secondary recipient mice at 4 weeks after transplantation ($n$=5; Student's $t$ test; * $P \leq 0.05$, ** $P \leq 0.001$, *** $P \leq 0.001$).

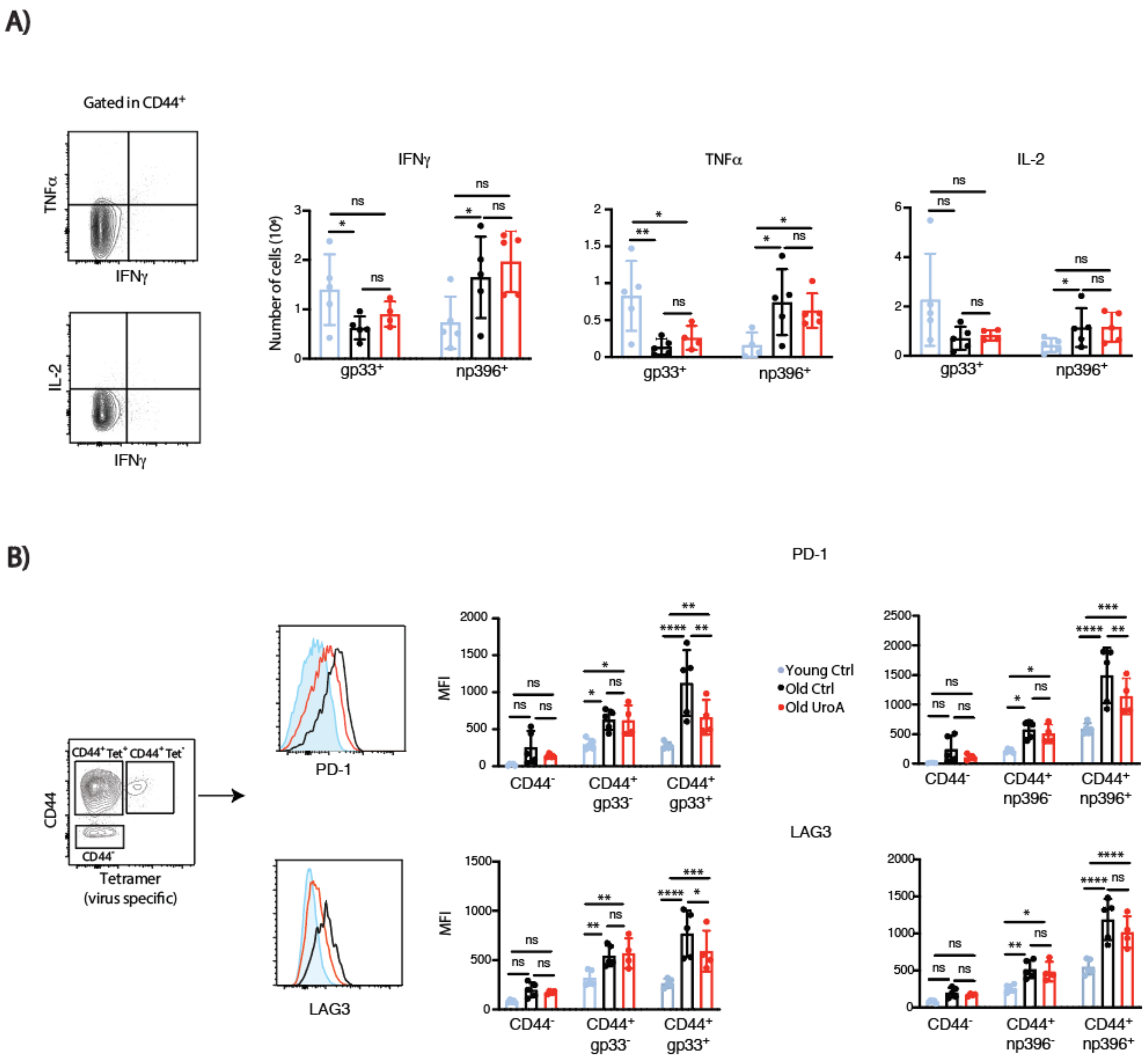


**Fig. S6. Analysis of inhibitory markers and cytokine expression.** (**A**) Expression of IFNγ, TNFα and IL-2 measured in CD44$^{+}$gp33$^{+}$ and CD44$^{+}$np396$^{+}$ CD8$^{+}$ T cells (**B**) Expression of co-co-inhibitory receptors PD-1 and Lag3 measured in non-activated (CD44-), non-virus specific activated (CD44$^{+}$gp33$^{-}$ and CD44$^{+}$np396$^{-}$) and virus-specific activated (CD44$^{+}$gp33$^{+}$ and CD44$^{+}$np396$^{+}$) CD8 T cells. (*n*=5; ANOVA; * P ≤ 0.05, ** P ≤ 0.001, *** P ≤ 0.001, **** P ≤ 0.0001).

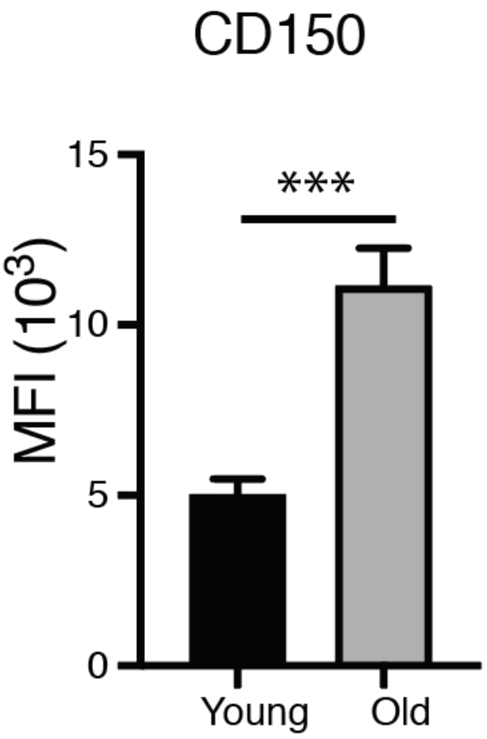


**Fig. S7. Old HSCs have higher CD150 expression.** Expression of CD150 measured by mean fluorescence intensity (MFI) in young and old HSCs (*n* = 4; Student's *t* test; * P ≤ 0.05, ** P ≤ 0.001, *** P ≤ 0.001).
